# WALLFLOWER, a RLK, simultaneously localizes to opposite sides of root hair cells & functions to position hairs

**DOI:** 10.1101/2023.03.16.533027

**Authors:** Jessica N. Toth, Cecilia Rodriguez-Furlan, Jaimie M. Van Norman

**Affiliations:** Department of Botany and Plant Sciences, Center for Plant Cell Biology, Institute of Integrative Genome Biology, University of California, Riverside, USA

**Keywords:** Arabidopsis root, root hair, epidermis, LRR-RLK, polar localization

## Abstract

Polarized cells are frequently partitioned into subdomains with unique features or functions. As plant cells are surrounded by walls, polarized cell shape and protein polarity in the plasma membrane are particularly important for normal physiology and development. We have identified WALLFLOWER (WFL), a transmembrane receptor kinase that is asymmetrically distributed at the inner face of epidermal cells and this localization is maintained independent of cell type. In epidermal hair (H) cells in the elongation and differentiation zones, WFL exhibits a dual polar localization, accumulating at the inner domain as well as at the root hair initiation domain (RHID). Furthermore, overexpression of WFL leads to a downward shift in root hair (RH) position suggesting WFL operates in a signaling pathway that functions across H cells to inform RH position. WFL asymmetric distribution and function is affected by deletion of the intracellular domains resulting in its mislocalization to the outer polar domain of H cells and exclusion from RHIDs and bulges. Thus, our results demonstrate that in epidermal H cells the WFL intracellular domains are required to direct its dual polar localization and influence RH position.

**ONE SENTENCE SUMMARY:** A receptor kinase with dual polar localization, to the inner polar domain and root hair initiation domain, in root epidermal cells, requires its intracellular domain for localization and function.

## INTRODUCTION

Eukaryotic cells are frequently partitioned into subdomains with unique features or functions and can therefore be described as polarized. Cell polarity can be defined as asymmetry in the localization of subcellular constituents and proteins and/or in cell morphology. One of the primary ways that cell polarity is achieved is through the preferential accumulation of proteins at specific subcellular locations. Partitioning of the plasma membrane (PM) can impact protein function, activity, and stability; therefore, it has tremendous regulatory potential in intercellular communication, development, and environmental interactions (Łangowski et al., 2016; Van Norman, 2016; Nakamura and Grebe, 2018). Because plant cells are fixed in place due to the cell wall, polarized protein accumulation is required for diverse physiological and developmental processes, such as asymmetric cell division, localized cell growth, long- and short-range signal transduction, and directional transport (Petrásek and Friml, 2009; Takano et al., 2010; Breda et al., 2017).

The Arabidopsis root is an excellent system to study various aspects of cell polarity and to link it to cellular or organ function and development. The root is composed of cells that are primarily cuboidal in shape and organized into concentric layers around the central vascular tissues with individual files of cells extending throughout the longitudinal axis (Dolan et al., 1993). In this axis, the root is divided into three developmental zones: the meristematic, elongation, and differentiation zones, within which cells divide, elongate, or mature, respectively (Benfey and Scheres, 2000). These organizational features allow for straightforward analysis of protein localization and growth, morphology, and developmental phenotypes in the root.

The root epidermis is composed of two cell types, hair (H) and nonhair cells (NH). Cells with H cell identity will form long, thin tubular extensions called root hairs (RHs), which are dramatic examples of morphological cell polarity. As RHs substantially increase root surface area, they are functionally important for efficient uptake of water and nutrients as well as plant anchorage (Grierson et al., 2014). Development and differentiation of the root epidermis and formation of RHs reveals links between subcellular polarity, tissue patterning, and structural polarity in terms of polarized cell growth. Additionally, RHs are an advantageous model to study how cellular polarity is achieved and maintained during development (Schiefelbein and Somerville, 1990).

The initiation of RHs requires the establishment of an additional polar domain within the context of the cell’s existing polarity. Confined to a very small region, the polar RH initiation domain (RHID) is located on the outer face near the rootward end of epidermal H cells. One of the first proteins to be positioned at the RHID is ROP GUANINE NUCLEOTIDE EXCHANGE FACTOR 3 (GEF3) followed by recruitment of RHO-RELATED PROTEIN FROM PLANTS 2 (ROP2) proteins (Denninger et al., 2019). Preceding H cell differentiation, GEF3 and ROP2 are uniformly distributed in the PM of these cells but as differentiation proceeds, these proteins accumulate at the RHID and are visualized as a disc-shaped area where the cell wall begins to soften. Subcellular and cell wall components are trafficked to the RHID, which leads to local formation of a bulge on the H cell surface that increases in size and becomes more defined (Grierson et al., 2014).

After the RHID is established, a dome-shaped outgrowth or bulge is formed leading to a series of events including cell wall acidification and loosening that facilitates the onset of RH elongation through a process known as tip growth (Gilroy and Jones, 2000). Bulge formation and tip growth occur through intensive polarized secretion of cellular and cell wall materials to this specific region of the cell. Many proteins show higher accumulation at the growing tip of root hairs; some of these proteins may be caught up in the default secretion scheme of RH elongation, but others, including many involved in signaling, are important for maintaining tip growth and cell wall integrity. One example is FERONIA (FER), a member of the *Catharanthus roseus* RECEPTOR-LIKE KINASE 1-LIKE (CrRLK1L) subfamily of putative cell wall sensors, which is involved in many developmental processes, including vacuolar expansion for cell elongation (Dünser et al., 2019). Additionally, FER shows polar accumulation in specific developmental contexts, but is otherwise nonpolar (Zhu et al., 2020). In RH formation, FER forms a complex with ROPs and GEFs and regulates tip growth through accumulation of reactive oxygen species (ROS) at the RH tip. Loss of function of either FER or GEFs reduces ROS accumulation, resulting in RHs that are shorter than wild type (WT) and have abnormally shaped tips (Duan et al., 2010; Huang et al., 2013). Mutants of another, CrRLK1 related receptor, ERULUS (ERU), have a similar RH phenotype. ERU-GFP preferentially accumulates at the tip of elongating RHs where it acts as a cell wall sensor that regulates cell wall elasticity through inhibition of pectin methylesterase activity (Kwon et al., 2018; Schoenaers et al., 2018). Thus, polarized proteins are essential for normal RH development, with cell wall sensing being crucial to maintain cell wall integrity and prevent cell rupture during the dramatic, local elongation of RHs.

Here we identify and characterize a polarly localized leucine-rich repeat receptor-like kinase (LRR-RLK) named WALLFLOWER (WFL). WFL is localized to the inner polar domain of epidermal cells in the elongation and differentiation zones and, in H cells, WFL maintains this polarity as it accumulates at the RHIDs and bulges at the outer polar domain of the PM. This unusual localization positions WFL simultaneously on opposite sides of H cells. This polar localization appears to be linked to function as WFL overexpression perturbs RH position in the longitudinal axis, leading to a downward shift. Furthermore, overexpression of WFL lacking the intracellular domain does not alter RH position indicating that signal transduction is related to this function. Our results also show that WFL polar accumulation and maintenance is mainly achieved by *de novo* protein synthesis and secretion through a Brefeldin A (BFA) dependent endomembrane trafficking pathway. WFL polarization appears to be determined by its intracellular domains as expression of truncated WFL is mislocalized to the outer polar domain of H cells and remarkably, fails to accumulate at RHIDs and bulges. Additionally, this truncated version of WFL is directed to different cellular domains in different cell types indicating that its intracellular portion is needed for correct, cell type-specific polar delivery. Given these results, we propose that polarly localized WFL participates in an epidermal signaling pathway that links cues from the root’s inner cell layers with polar growth at the outer epidermal surface, informing RH position.

## RESULTS

### *WFL* is expressed primarily in LRC and epidermal cells

We identified WALLFLOWER (WFL), encoded by At5g24100, as an LRR-RLK putatively involved in signaling and RH development based on its predominant expression in H cells in the elongation and differentiation zones (Brady et al., 2007; Li et al., 2016). To validate these data *in planta*, we drove expression of endoplasmic reticulum-localized green fluorescent protein (erGFP) with the putative *WFL* promoter (*pWFL*) in WT seedlings. *pWFL* activity was observed in the lateral root cap (LRC), but was not detectable in other cell types in root meristem (Figure 1D-E). Consistent with the expression data, in elongation and differentiation zones, *pWFL* activity was detected in epidermal cells with preferential activity in H cells. Additionally, we detected *pWFL* activity in pericycle cells along with weaker activity in cortex cells (Figure 1B-C).

**Figure 1.**
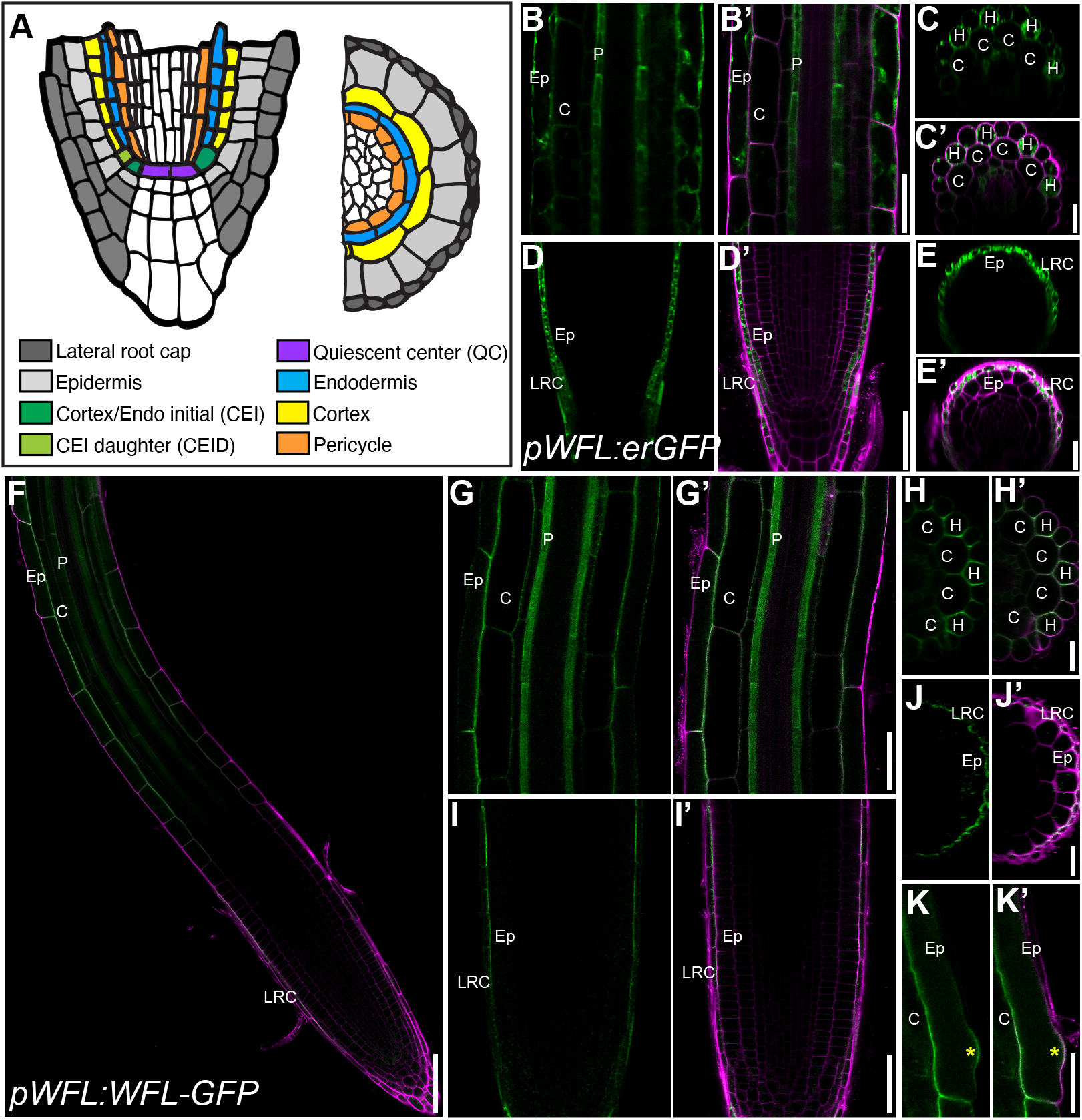
*pWFL* is active in the lateral root cap and epidermis and WFL-GFP localizes to the inner polar domain of these cell types. (A) Schematic representation of cell types in the Arabidopsis root in longitudinal and transverse views. (B-K) Confocal images of WT roots expressing (B-E) *pWFL:erGFP* and (F-K) *pWFL:WFL-GFP* and stained with propidium iodide (PI) to show cell outlines. Adjacent panels show GFP alone (α) and GFP + PI merged (α’), except in (F) where only the merged image is shown. (B and C) In the elongation and differentiation zones, *pWFL* is most active in pericycle and epidermal cells with (C) higher activity in H cells compared to NH cells. (D and E) In the meristematic zone, *pWFL* is active in the cell layers of the LRC. (F) WFL*-*GFP localization in the root tip. (G and H) In the elongation and differentiation zones, WFL-GFP localizes to the inner polar domain of (G) epidermal cells with (H) preferential accumulation in H cells. (K) WFL-GFP also localizes to RH bulges (yellow asterisk). (I and J) WFL-GFP localizes to the inner polar domain of the outermost layer of the LRC. Abbreviations: LRC, lateral root cap; Ep, epidermis; C, cortex; P, pericycle; H, hair cell. Scale bars: 50 μm in (F); 25 μm in (B, D, G, and I); 10 μm in all others.

### WFL accumulates asymmetrically at the PM

To examine WFL protein accumulation in Arabidopsis roots, we generated a GFP fusion under control of *pWFL* (*pWFL:WFL-GFP*). In the meristematic zone of WT roots expressing *pWFL:WFL-GFP*, we detect the protein only in the outermost cell layer of the LRC (Figure 1F, 1I-J). Consistent with the observed promoter activity, in the elongation and differentiation zones WFL-GFP accumulates in cells of the epidermis, cortex, and pericycle. In the LRC and epidermis, WFL-GFP is polarly localized to the inner polar domain of the PM (Figure 1F-J). Interestingly, in the epidermis, WFL has higher accumulation in H cells and is also localized to the RH bulge (Figure 1K). In cortex cells, we were unable to determine whether WFL-GFP was polarly localized due to the fluorescent signal from the adjacent epidermis. Also, in the pericycle, WFL-GFP signal appears diffuse, making it difficult to assess how WFL is distributed at the PM (Figure 1F-G). These results indicate that WFL is asymmetrically distributed along the PM in LRC and epidermal cells at different stages of differentiation.

### WFL-GFP localizes to the inner polar domain regardless of cell type

With the endogenously expressed reporter, it was difficult to assess the polar distribution of WFL-GFP within internal cell layers of the root. To address this, we misexpressed WFL-GFP using cell type- and tissue-specific promoters. WFL-GFP expressed under the control of the *WEREWOLF* promoter (*pWER, (Lee and Schiefelbein, 1999)*, which is specifically expressed in the epidermis and LRC (Figure 2A), confirmed WFL localization to the inner polar domain of these cell types (Figure 2B-C). We also misexpressed WFL-GFP in the endodermis, cortex/endodermal initial (CEI), cortex/endodermal initial daughter (CEID), and quiescent center (QC) (Figure 2D) using the *SCARECROW* promoter (*pSCR, (Wysocka-Diller et al*., *2000; Levesque et al*., *2006)*. We found that WFL-GFP localized towards the stele in these cell types, accumulating at the inner polar domain of endodermal and initial cells and the shootward polar domain in the QC (Figure 2E). Additionally, we misexpressed WFL-GFP in immature and mature cortex cells using the promoters of *CORTEX2* (*pCO2, (Heidstra et al*., *2004; Paquette and Benfey, 2005)*) and *CORTEX* (*pC1, (Lee et al*., *2006)*), respectively. We were unable to detect any GFP signal in immature cortex cells (not shown) but observed that WFL-GFP localized to the inner polar domain of mature cortex cells (Figure 2G). Altogether, our results show that polar localization of WFL-GFP is oriented inwards, towards the stele in all cell types examined. This suggests that, as proposed for some nutrient transporters (Alassimone et al., 2010), WFL localization to the inner polar domain may be informed by a cue originating from the stele.

**Figure 2.**
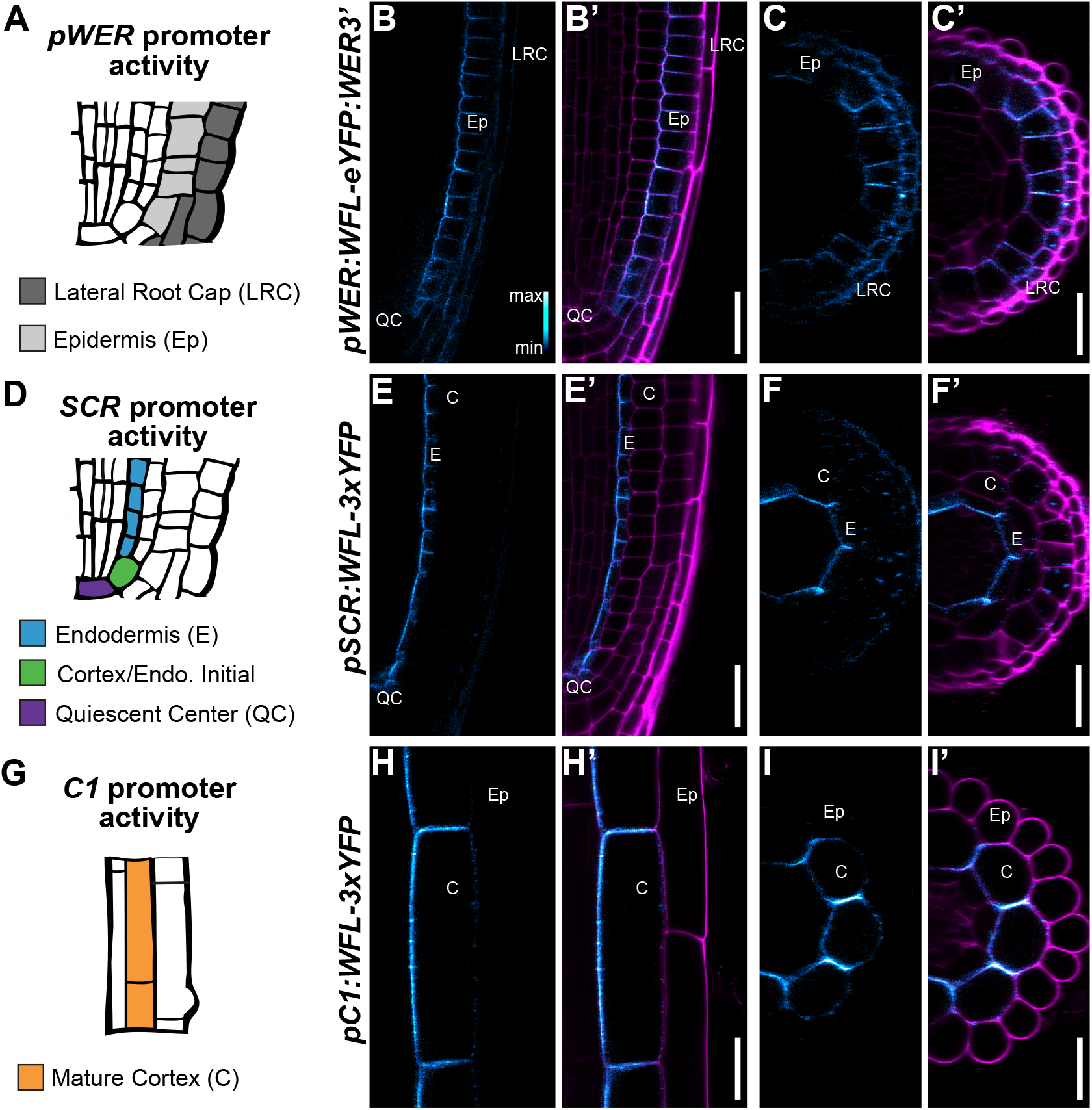
WFL-GFP localizes to the inner polar domain regardless of cell type. (A, D, and G) Schematics indicating activity of various promoters in specific cell layers. (A) *pWER* is active in LRC and epidermis. (D) *pSCR* is active in endodermis, CEI and QC. (G) *pC1* is active in mature cortex cells. (B, C, E, F, H, and I) Confocal images in longitudinal (B, E, and H) and transverse (C, F, I) planes of WT roots expressing WFL-GFP/YFP driven by cell layer-specific promoters (*pWER, pSCR*, and *pC1*) and stained with propidium iodide (PI) with adjacent panels showing GFP alone (false colored to show signal intensity, (α) and GFP + PI merged (α’). (B-C) In the LRC and epidermis, WFL-eYFP localizes to the inner polar domain. (E-F) In endodermis and (H-I) the mature cortex, WFL-GFP localizes to the inner polar domain. Abbreviations: LRC, lateral root cap; Ep, epidermis; C, cortex; E, endodermis; QC, quiescent center. Scale bars: 25 μm.

### WFL-GFP is dynamically trafficked to and from the PM

Proteins with polar localization can be directed to the PM by targeted secretion and/or maintained at the PM by endocytosis and recycling (Rodriguez-Furlan et al., 2019; Raggi et al., 2020). Beyond its PM localization, in growing RHs WFL-GFP is detected in mobile intracellular compartments that most likely correspond to highly dynamic endomembrane traffic (Movie SM1). To understand how endomembrane trafficking contributes to the polar distribution of WFL-GFP at the PM, we performed a series of chemical treatments on roots expressing *pWFL* driven WFL-GFP (Figure 3A). Treatments with Brefeldin A (BFA), an inhibitor of Golgi trafficking that affects secretion to the PM, generated intracellular accumulations consistent with BFA bodies. These accumulations indicate that WFL-GFP is trafficked to the PM via a BFA-sensitive mechanism (Figure 3B and F).

**Figure 3.**
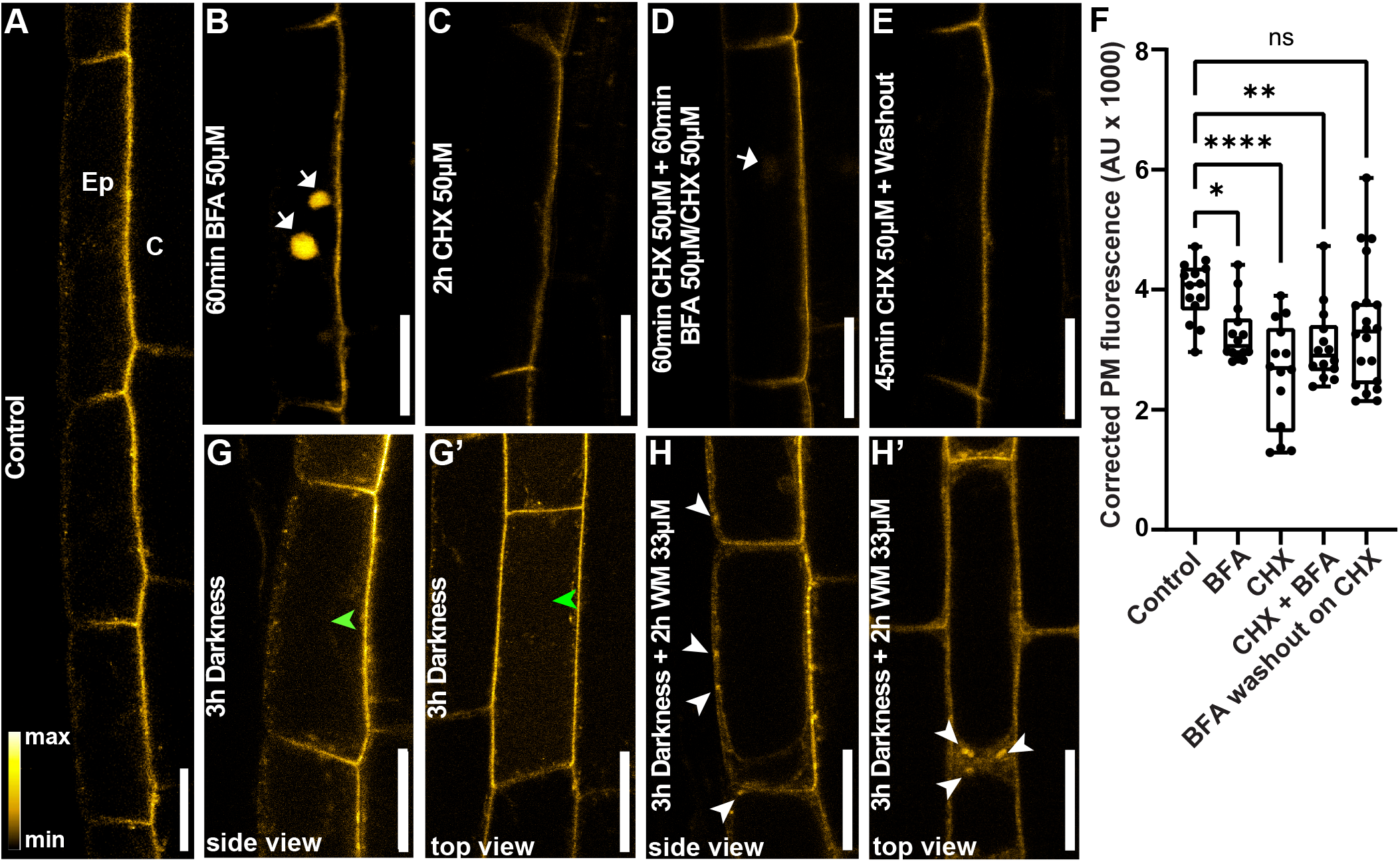
WFL-GFP is dynamically localized at the PM and trafficked to the vacuole for degradation. (A-F) Confocal images of unstained WT roots expressing *pWFL:WFL-GFP*. (A) In untreated control, WFL-GFP localizes to the inner polar domain of epidermal cells. (B) 60-minute BFA treatment results in WFL-GFP accumulation in BFA bodies. (C) 2-hour CHX treatment reduces WFL-GFP fluorescence at PM. (D) WFL-GFP weakly accumulates in BFA bodies upon cotreatment with CHX and BFA. White arrows indicate WFL-GFP in BFA bodies. (E) 2-hour CHX treatment followed by BFA washout does not result in appreciable signal recovery at the plasma membrane. (F) Graph shows the quantification of WFL corrected fluorescence intensity in arbitrary units (AU) at the PM after the indicated treatments. Data shown are representative results of experiments with at least three independent replicates. Bars indicate min. to max. values and 1-4 stars indicate statistical significance (P values ≤0.05, one-way ANOVA using Dunn’s multiple comparison test). (G and G’) 3-hour dark treatment induces WFL-GFP trafficking to vacuole. Green arrowheads indicate WFL-GFP accumulation in the vacuole lumen. (H and H’) 2-hour Wortmannin (Wm) treatment results in accumulation of WFL-GFP in Wm bodies and inhibition of vacuolar trafficking. (G and H) Side view and (G’ and H’) top view of epidermal cells. White arrowheads indicate WFL-GFP in Wm bodies. Abbreviations: Ep, epidermis; C, cortex. Scale bars: 20 μm.

WFL-GFP accumulation into BFA bodies could be attributed solely to secretion of newly synthesized WFL-GFP or to protein returning to the PM after endocytosis and recycling to maintain the polarized pool of proteins. To investigate this, we first treated roots with cycloheximide (CHX), an inhibitor of protein synthesis and after 2 hours we observed a considerable reduction in WFL-GFP signal at the PM indicating a high rate of *de novo* protein secretion and turnover (Figure 3C and F). We next pre-treated the roots with CHX for 60 minutes and added BFA and incubated for an additional 60 minutes. After the co-treatment, WFL accumulation in BFA bodies was nearly abolished, with only a faint signal remaining detectable (Figure 3D and F). Therefore, the majority of the signal observed in BFA bodies can be attributed to the endomembrane trafficking of newly synthesized WFL-GFP. Furthermore, after a 60-minute BFA treatment followed by wash out in the presence of CHX, WFL-GFP signal at the PM is recovered to values similar to the control (Figure 3E and F); again, indicating a high rate of protein turnover.

As our results indicate a high rate of protein turnover, we explored whether WFL is actively degraded by a Wortmannin (Wm)-sensitive pathway. Wm is an inhibitor of phosphoinositide synthesis that has been reported to inhibit endocytic trafficking of PM proteins towards the vacuole. Additionally, it has been shown that darkness induces internalization and trafficking of PM proteins to the vacuole and changes vacuolar pH, which delays degradation allowing fluorescent protein detection at the vacuole lumen (Kleine-Vehn et al., 2008). *pWFL:WFL-GFP* expressing roots were exposed to a 3 hour dark treatment to increase WFL-GFP transport to the vacuole evidenced by the GFP detection at the lumen in the epidermal cells (Figure 3G and G’’). Upon a 2-hour treatment with Wm in dark conditions, we observed characteristic doughnut-shaped intracellular accumulations of WFL-GFP and a considerable decrease in fluorescent signal at the vacuole lumen (Figure 3H and 3H’). These results indicate that WFL-GFP is actively endocytosed and trafficked to the vacuole by a Wm sensitive pathway.

### The WFL kinase domain is necessary for its polar distribution

To determine whether specific protein domains inform WFL polar localization at the PM, we created a truncated version by removing the intracellular region, which consists of the juxtamembrane (Jx) and kinase (K) domains, and fused this truncation to GFP under *pWFL* (*pWFL:WFLΔJxK-GFP*). Similar to full-length WFL-GFP, we detected accumulation of WFLΔJxK-GFP in LRC cells of the meristematic zone, as well as in epidermal, cortex, and pericycle cells in the elongation and differentiation zones (Figure 4A-G). However, WFL localization to the inner polar domain and the RHID was strongly impacted. Indeed, WFLΔJxK-GFP polar localization appears to switch from the inner to the outer polar domain in LRC and epidermal NH and H cells (Figure 4A-G) and is specifically excluded from the RHID and bulge of H cells (Figure 4E-F). Similar results were obtained by removing only the kinase domain (*pWFL:WFLΔK-GFP*, Figure S1). When compared to WFLΔJxK-GFP, the WFLΔK-GFP signal is lower, suggesting a reduction in secretion to the PM and/or protein instability. These results suggest WFL localization is highly regulated and that the intracellular domains are required for normal WFL polar localization.

**Figure 4.**
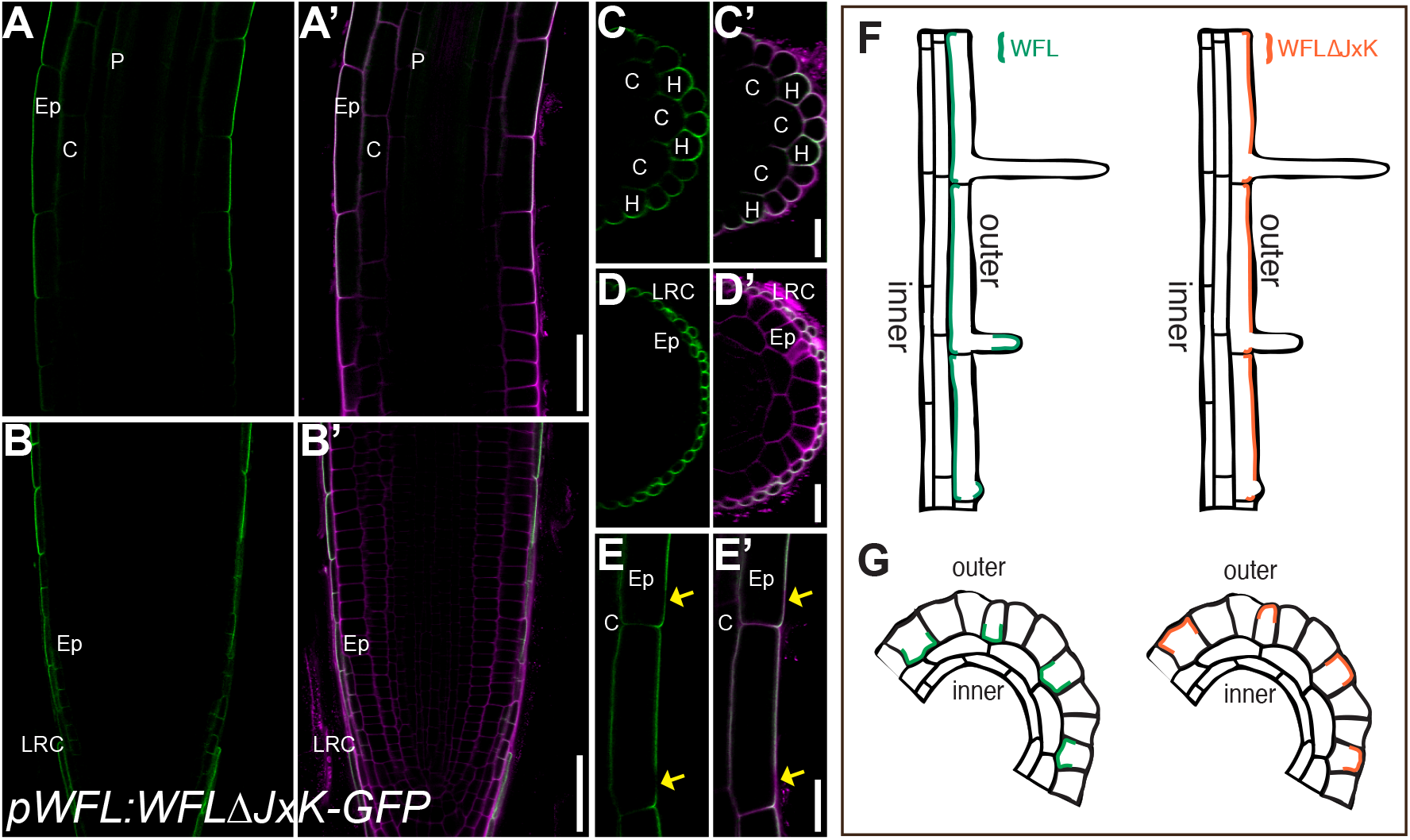
Deletion of the intracellular domains redirects WFL localization. (A-E) Confocal images of WT roots expressing WFLΔJxK-GFP driven by *pWFL* (*pWFL:WFLΔJxK-GFP)* and stained with propidium iodide (PI) to show cell outlines. Adjacent panels show GFP alone (α) and GFP + PI merged (α’). (A and C) In elongation and differentiation zones, WFLΔJxK-GFP localizes to the outer polar domain of epidermal cells with (C) preferential accumulation in H cells. (E) WFLΔJxK-GFP is excluded from RHIDs (yellow arrows). (B and E) WFLΔJxK-GFP localizes to the outer polar domain of the outermost cell layer of the LRC. (F and G) Schematics of WFL-GFP and WFLΔJxK-GFP localization in median (F) longitudinal and (G) transverse views. Abbreviations: LRC, lateral root cap; Ep, epidermis; C, cortex; P, pericycle; H, hair cell. Scale bars: 25 μm in (A, B, and E); 10 μm in all others.

### WFL intracellular domains are important for cell type-specific polar localization

WFL appears to be oriented by a stele-derived cue that coordinates its inner polar distribution in different cell types. Therefore, we explored whether deleting the WFL cytoplasmic domain alters the localization of WFL in different cell types. When expressed from *pWER* and *pCO2*, WFLΔJxK-3xYFP preferentially localizes to the outer polar domain of LRC/epidermal and immature cortex cells near the QC (Figure 5A-B and E), respectively. Upon expression in mature cortex from *pC1*, WFLΔJxK-3xYFP was predominantly localized to the outer polar domain, however, some elongating cells showed nonpolar distribution of the protein (Figure 5C-D). Notably, when WFLΔJxK-3xYFP was expressed from *pSCR* there was no detectable signal in the primary root, however, in the endodermis and ground tissue stem cells of lateral roots, signal was detectable and showed a nonpolar distribution along the PM (Figure 5F). Thus, in contrast to full length WFL-GFP, there is no uniform interpretation of cues to polarize truncated WFL among the different cell types examined. These data indicate that WFL localization is highly regulated and its polarization towards the stele requires the intracellular domains, without which truncated WFL is misdirected in different cell types and developmental contexts.

**Figure 5.**
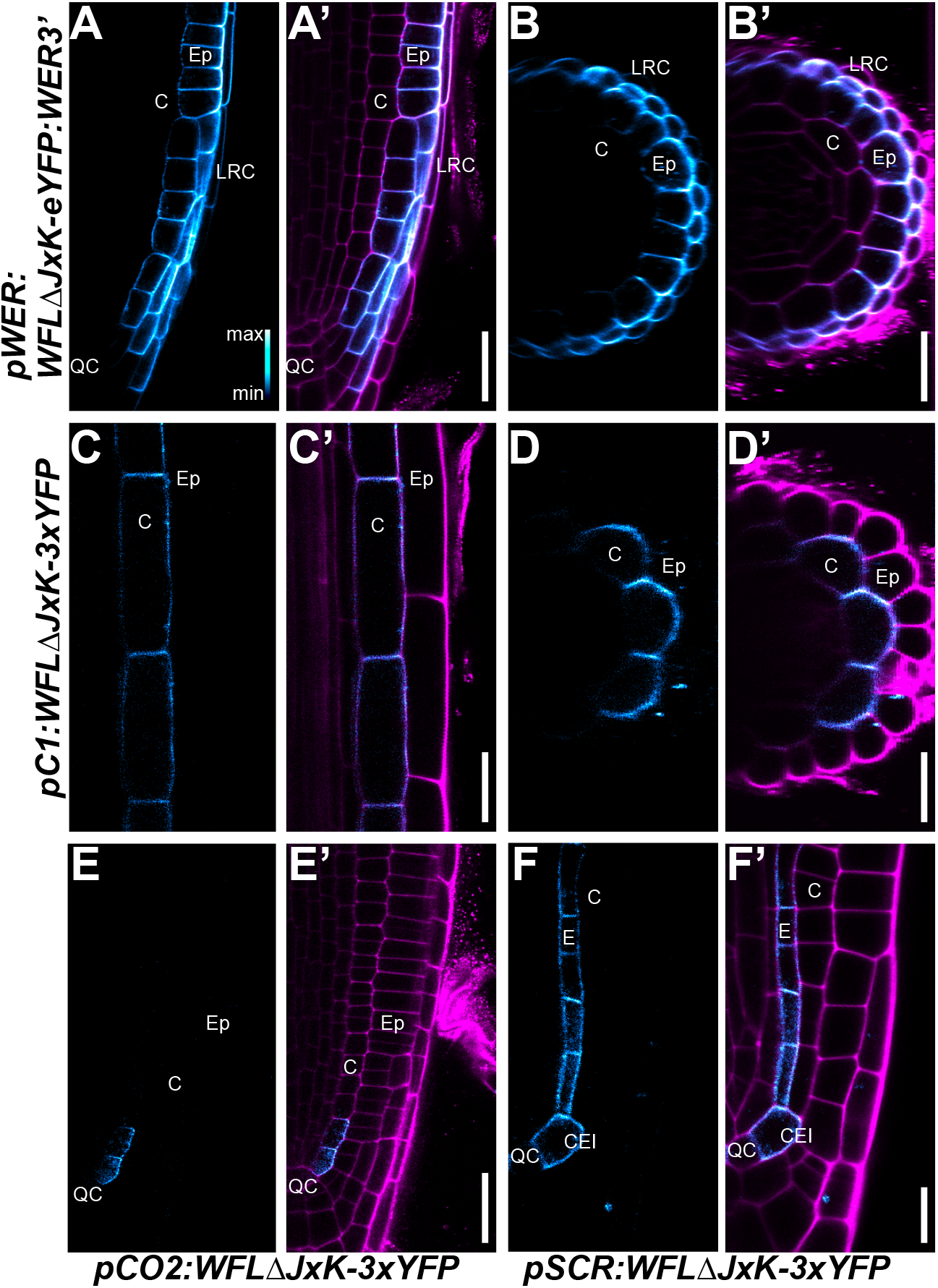
Truncated WFL predominantly localizes to the outer polar domain or is nonpolar. (A-D) Confocal images of WT roots expressing WFLΔJxK-eYFP/3xYFP driven by cell layer-specific promoters (*pWER, pCO2, pC1*, and *pSCR*) and stained with propidium iodide (PI) with adjacent panels showing YFP alone (false colored to show signal intensity, (α) and YFP + PI merged (α’) (A, C, E, F) show images in longitudinal planes and (B, D. E, and F) in the transverse planes (A-B) In the LRC and epidermis, WFLΔJxK-eYFP localizes to the outer polar domain. (C-D) In mature cortex cells, WFLΔ JxK-3xYFP preferentially localizes to the outer polar domain. (E) In immature cortex cells, WFLΔJxK-3xYFP localizes to the outer polar domain. (F) In ground tissue initials and endodermal cells of lateral roots, WFLΔJxK-3xYFP is nonpolar. Abbreviations: LRC, lateral root cap; Ep, epidermis; C, cortex; E, endodermis; CEI, cortex/endodermal initial; QC, quiescent center. Scale bars: 25 μm.

### *WFLΔJxK* is polarly distributed but actively excluded from WFL accumulation domains

To further characterize the opposite localization of WFL and WFLΔJxK in H cells, we closely followed their respective distributions during the different stages of RH development. WFL-GFP is present at the inner polar domain of elongating H cells and gradually appears at the RHID and is present at the bulge. In contrast, WFLΔJxK-GFP is present at the outer polar domain in elongating H cells and gradually decreases its accumulation at RHIDs and later at the bulge (Figure 6A-B). Additionally, WFL-GFP shows the highest fluorescence intensity at the center of RHIDs and RH bulges (Figure 6C-F), whereas roots expressing *pWFL:WFLΔJxK-GFP* exhibited lower fluorescence intensity at the center of RHIDs and RH bulges with higher fluorescence above and below the developing RH (Figure 6G-J). Together, these results indicate that WFL intracellular domains are important for polarized accumulation of WFL at specific domains of the PM.

**Figure 6.**
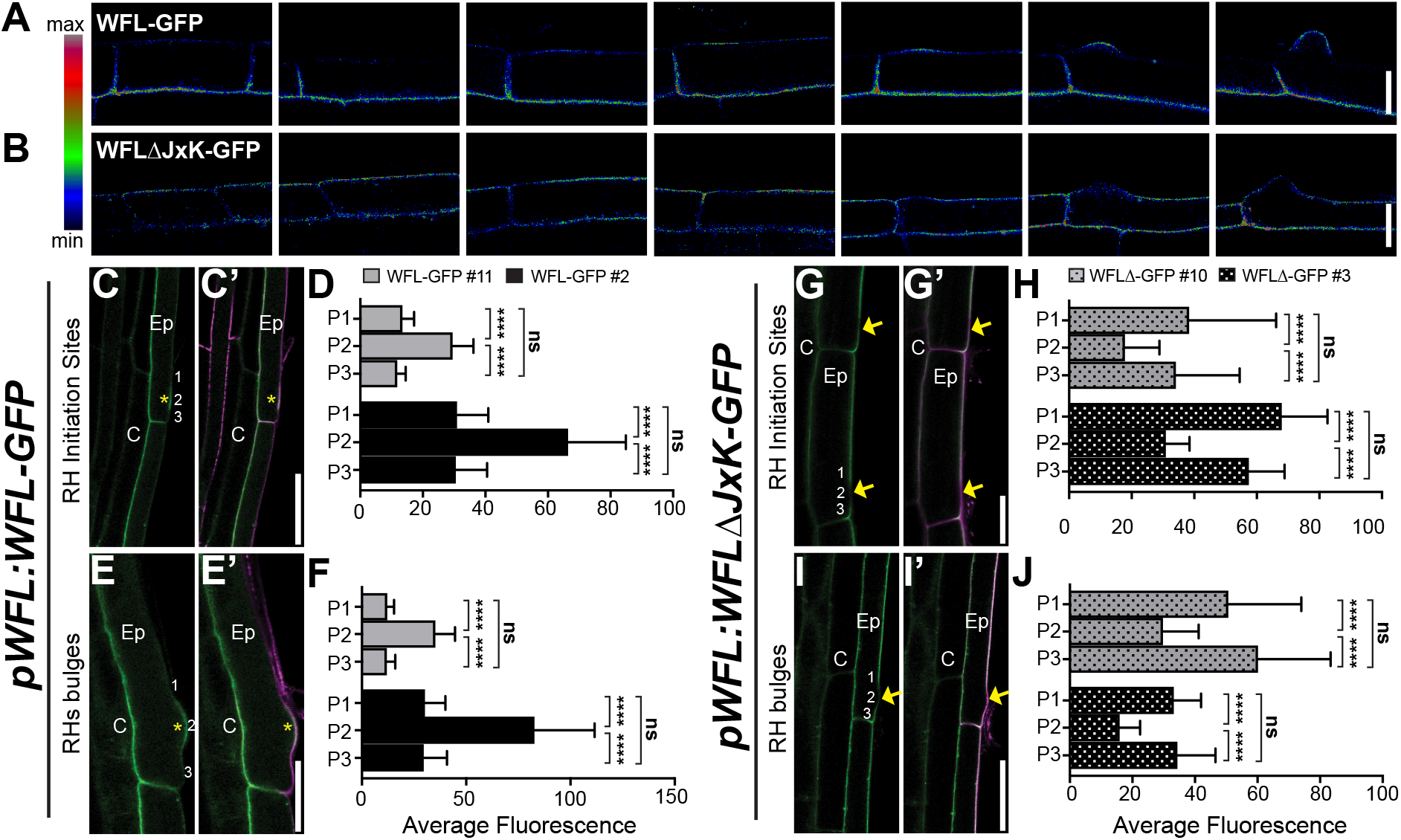
WFL-GFP and WFLΔJxK-GFP have reciprocal localization at RHIDs and bulges. (A and B) Confocal images of WT root H cells expressing *pWFL* driven (A) WFL-GFP and (B) WFLΔJxK-GFP (GFP false colored to show signal intensity). As H cell development progresses, (A) WFL-GFP localizes to RHIDs and bulges while (B) WFLΔJxK-GFP is excluded from these sites. (C, E, G, and I) Confocal images of WT roots expressing *pWFL* driven WFL-GFP and WFLΔJxK-GFP and stained with propidium iodide (PI) with adjacent panels showing GFP alone (α) and GFP + PI merged (α’). (C and E) WFL-GFP localizes to the inner polar domain of H cells and to (C) RHIDs as well as (C) bulges. (D and F) Quantification of fluorescence intensity at 3 positions - above, at the center, and below of (D) RHIDs and (F) bulges in two independent transgenic lines. (G and I) WFLΔJxK-GFP localizes to the outer polar domain of epidermal cells and is excluded from (G) RHIDs and (I) bulges. (H and J) Quantification of fluorescence intensity above, below, and at (H) RHIDs and (I) bulges in two independent transgenic lines. For graphs: error bars, SD; student’s t test, **** p<0.0001. Two different transgenic lines for each reporter were used as biological replicates, with n= 15-20 roots and 2-3 cells per root per replicate. Yellow asterisks and arrows indicate RHIDs and bulges for WFL-GFP and WFLΔ JxK-GFP, respectively. Numbers indicate positions of fluorescence intensity measurements relative to RH apex, 1= above, 2= center, 3= below. Abbreviations: Ep, epidermis; C, cortex. Scale bars: 20 μm in (A and B) 50 μm in (C and I); 25 μm in (E and G).

### Overexpression of WFL affects the position of RHIDs and bulges

In roots overexpressing WFL-GFP, we observed that RH position is perturbed. In WT plants the RHID is normally located approximately 10 µm from the rootward edge of epidermal H cells (Figure 7A, (Grierson et al., 2014). However, in *pWFL:WFL-GFP* roots, we observed that bulge formation is consistently shifted downward toward the rootward edge of H cells (Figure 7B). To quantify this phenotype, we classified RH bulge position as WT (normal) or shifted downward, where shifted RHs have no measurable distance between the RH bulge and the rootward edge of the cell. We used confocal microscopy to visualize bulge position in two independent *pWFL:WFL-GFP* transgenic lines and concluded that bulge position is indeed shifted towards the rootward edge of H cells in these roots (Figure 7D). These results indicate that overexpression of WFL-GFP in a WT background leads to a defect in RH positioning.

**Figure 7.**
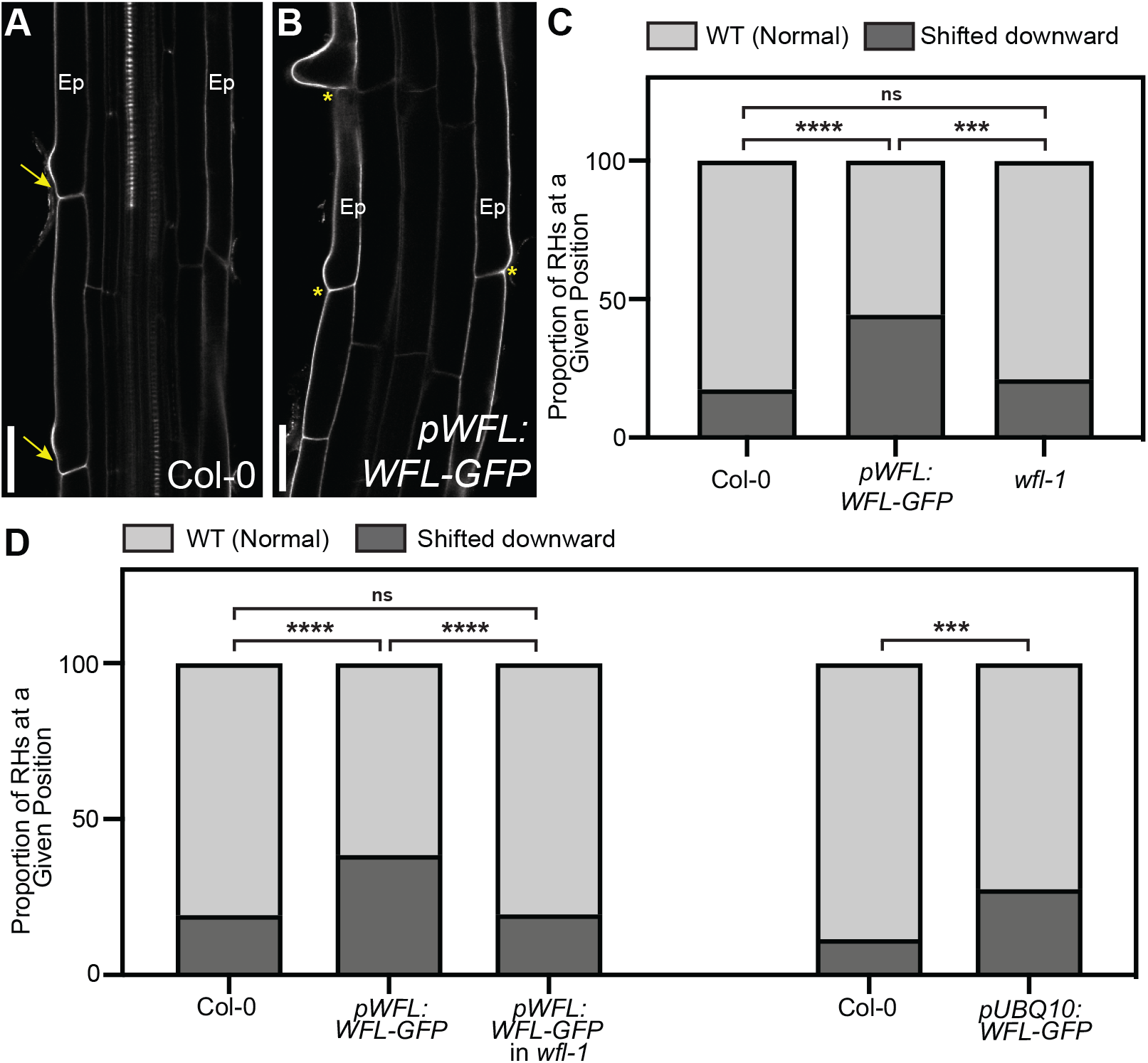
Overexpression of WFL-GFP shifts root hair position downward toward the rootward edge of hair cells. (A and B) Confocal images showing propidium iodide (PI) stain (gray) to show cell outlines of (A) WT and (B) WT roots expressing *pWFL:WFL-GFP*. (A) RHs in WT roots have a defined space between the rootward edge of H cells and the site of the RH bulge (yellow arrows). (B) In contrast, in roots expressing *pWFL:WFL-GFP*, RHs are shifted downward towards the rootward edge of H cells (yellow asterisks). (C and D) RHs were binned into 2 categories based on position and quantified according to genotype. (C) RH bulge position in *wfl-1* and WT is the same, whereas RH bulges are shifted downwards when WFL-GFP is overexpressed by either *pWFL* or *pUBQ10*. No change in RH bulge position is observed when *pWFL:WFL-GFP* is expressed in *wfl-1*. For graphs: student’s t test, *** p<0.001 and **** p<0.0001. Data shown is from one (of two) independent transgenic lines per reporter (with similar results for each line in each replicate) and 3-4 biological replicates combined, with n=15 roots and 3-5 cells per root per replicate. Abbreviations: Ep, Epidermis. Scale bars: 25 μm.

Interestingly, during our examination of *pWFL:WFL-GFP* in the WT (Col-0) background we observed that these roots appeared damaged more frequently than nontransgenic WT or *pWFL:erGFP* (in WT) roots, as evidenced by the presence of propidium iodide (PI) within cells. Incidents of damage were observed in >4 independent transgenic lines and by separate researchers (not shown). We hypothesized that these plants were sensitive to being mounted on slides, a type of mechanical stress, possibly due to a defect in cell wall integrity. To test this we transferred seedlings expressing *pWFL:WFL-GFP* from our standard growth medium to medium deficient in phosphate (-Pi), which has been reported to rigidify the walls of elongating root cells (Péret et al., 2011; Balzergue et al., 2017). After a 24-hour treatment on -Pi media, these roots showed substantially less damage (PI penetration) and WFL-GFP localization was unaffected. This suggests that expression of WFL-GFP in a WT background has additional phenotypic consequences. Together with the shift in RH position, it is tempting to propose that WFL is involved in cell wall modification and that this activity along with its polarized presence at the RHID and bulge might alter RH positioning upon overexpression.

To confirm the level of *WFL* overexpression in *pWFL:WFL-GFP* transgenic plants compared to non-transgenic WT controls, we conducted RT-qPCR. *WFL* transcript levels were approximately 2-fold higher in transgenic plants compared to WT and were similar between the two independent transgenic lines (Figure S2B). Similarly, when bulge position was quantified in another overexpression line where WFL-GFP is expressed from the constitutively active *UBIQUITIN10* promoter (*pUBQ10, pUBQ10:WFL-GFP*) we observed that bulges were shifted downward (Figure 7D). Notably, the proportion of shifted bulges in *pUBQ10:WFL-GFP* roots was similar to that of *pWFL:WFL-GFP* roots confirming the relationship between WFL overexpression and altered RH positioning. These results suggest overexpression (above the endogenous level) or increased copy number of WFL leads to a shift in RH position, however, an alternative explanation is that the GFP fusion somehow interferes with WFL function causing this phenotype. To test this, we generated an untagged version of the transgene (*pWFL:WFL*) and upon its expression in WT, we observed a similar shift in RH bulge position as seen in WFL-GFP fusions (Figure S2A). Thus, the fusion of GFP to WFL cannot explain the abnormal RH bulge position phenotype, suggesting expression level or increased copy number leads to this phenotype.

To further characterize WFL function, we generated a putative null allele, *wfl-1*, with reduced *WFL* expression (Figure S3), however, we did not observe any RH positioning phenotype. Additionally, expression of *pWFL:WFL-GFP* in the *wfl-1* background did not result in a shift in the position of RH bulges (Figure 7D) indicating a reduction in expression or functional copy number alleviates this phenotype. Finally, we examined RH bulge position in WT and *wfl-1* roots expressing *pWFL:WFLΔJxK-GFP* and found that bulge position was unaffected in either genotype (Figure S2C). Taken together, these results indicate that only overexpression of full length WFL elicits a shift in RH bulges toward the rootward edge of H cells indicating that the WFL intracellular domains and/or subcellular localization are required for this phenotype.

## DISCUSSION

Polarization of proteins at the PM often precedes polarized cellular morphology. WFL polarity at the inner polar domain in epidermal cells during cell elongation and then its appearance at the RHID site implicate it in epidermal cell differentiation and RH positioning. The dual polarization of WFL is likely informed both by features of the WFL intracellular domain and cell type-specific factors. The importance of the WFL intracellular domains is highlighted by the change in WFL localization from the inner to the outer polar domain, upon their removal. This is further underscored by the reduced accumulation of WFLΔJxK-GFP at RHIDs and bulges. Our results suggest that polar localization of WFL is subject to precise spatiotemporal regulation. It is also clear that WFL intracellular domains are functionally important for RH positioning, as only overexpression of the full-length protein results in a shift in RH position. Thus, we propose that WFL functions in a signaling pathway that links cues from the inner cell layers of the root with polar growth at the epidermal surface to inform RH position..

Modification of cellular infrastructure is required to accommodate the drastic change in cellular polarity and morphology as a RH develops; therefore, there are multiple factors that can influence RH position. In the *deformed root hairs 1* (*der1*) mutant, in which *ACTIN2* is mutated, RHs often emerge in the middle of H cells, indicating that the cytoskeleton is important for normal RH positioning and bulge formation (Ringli et al., 2002). Furthermore, hormones also have a role in determining RH position, for example, positioning is affected in auxin/ethylene perception and biosynthesis mutants (Masucci and Schiefelbein, 1994a; Masucci and Schiefelbein, 1996).

One of the greatest obstacles during RH development is the cell wall, which must be modified to allow asymmetric, polarized growth of the RH. In *procuste1* (*prc1-1*) mutants, which have a defect in the *CELLULOSE SYNTHASE6* (*CESA6*) gene, RH position is shifted toward the rootward edge of H cells (Singh et al., 2008), similar to the phenotype observed in roots overexpressing WFL-GFP. In addition to shifted RHs, roots overexpressing WFL-GFP also appear to be sensitive to mechanical stress and this sensitivity can be alleviated by exposure to -Pi growth conditions, which rigidifies root cell walls. Intriguingly, WFL localizes to areas of the root where dramatic changes to the cell wall are taking place, including the outermost layer of the LRC, cells of the elongation zone, and RHIDs and bulges. The presence of WFL at these positions of cell wall modification together with a shifted RH position upon WFL overexpression make it very tempting to speculate that WFL is involved in sensing cell wall status and/or cell wall modification.

In further support of a role for WFL in cell wall sensing or modification, the Search Tool for the Retrieval of Interacting Genes/Proteins (STRING) database predicts interactions with a RH-specific proline-rich extensin-like family protein (EXT15, AT1G23720) and two RH-specific Class III (CIII) peroxidase (PRX) proteins (PRX27, AT3G01190 and PRX57, AT5G17820) (Szklarczyk et al., 2019). *prx57* mutants have shorter RHs with frequent bursting, indicating that PRX57 plays a role in cell wall modification during RH elongation (Kwon et al., 2015). It is known that LRR-RLKs interact with extensins, for example, FERONIA interacts with LEUCINE-RICH REPEAT/EXTENSIN 1 (LRX1) to coordinate cell wall loosening with cell elongation (Dünser et al., 2019). Further research into a possible role for WFL in cell wall sensing or modification through these cell wall-associated proteins and their potential interaction with WFL are intriguing areas for future study.

Endomembrane protein trafficking is closely related to the establishment and maintenance of PM protein polarity (Muroyama and Bergmann, 2019; Rodriguez-Furlan et al., 2019; Raggi et al., 2020). Our results indicate that WFL polarity at the PM is primarily maintained by constant secretion and degradation. The WFL biosynthetic secretory traffic is directed through a BFA sensitive pathway, however, BFA interference with protein delivery does not alter WFL polarity as it has been described for PIN1 (Tanaka et al., 2014). Additionally, endocytosis and recycling does not appear to be responsible for maintaining the pool of WFL at the PM, which instead relies mainly on *de novo* protein synthesis. Therefore, WFL polar secretion appears to be similar to that of other laterally localized proteins, such as POLAR AUXIN TRANSPORT INHIBITOR-SENSITIVE 1/PLEIOTROPIC DRUG RESISTANCE 9 (PIS1/PDR9/ABCG37) (Langowski et al., 2010; Łangowski et al., 2016).

While polar localization of full-length WFL is likely informed by organ-level polarity cues, removal of the intracellular domains indicates that the kinase domain is essential for different cells to interpret these cues and localize WFL. Unlike WFL-GFP, misexpression of WFLΔJxK-GFP reveals differential localization depending on cell identity and developmental context. WFLΔJxK-GFP localizes to the outer polar domain of epidermal cells and the immature cortex cells, whereas in mature cortex cells, WFLΔJxK-GFP is also sometimes present at the inner PM domain. Strikingly, WFLΔJxK-GFP cannot be detected in endodermal cells of the primary root, but in lateral roots, exhibits nonpolar localization in the endodermis. Therefore, it is possible that when the intracellular domains are absent, the protein is redirected to different secretion pathways in different cell types. This hypothesis is consistent with the existence of multiple endomembrane trafficking pathways governing polar localization of transmembrane receptors in plants (Li et al., 2017). These results underscore the necessity of WFL intracellular domains to direct its polar localization, which is informed by context specific factors that take into account cell type and developmental stage.

WFLΔJxK-GFP is secreted to H cell domains where full length WFL does not normally accumulate; indeed, WFLΔJxK-GFP appears to be excluded from the inner domain and from the RHID. The distribution of these two proteins is particularly intriguing and suggests that their secretion to the PM is subject to strict regulation. It is unclear how this contrasting localization is achieved, but it could be explained by interaction, or lack thereof, with other proteins. Specifically, interaction of the WFL cytoplasmic domain with another protein that is polarly localized at the RHID but excluded from the rest of the outer polar domain could explain WFL polar localization. Recently, polar localization of GEF3 to the RHID was shown to be necessary for ROP2 recruitment to this site (Denninger et al., 2019). Therefore, it is possible that in the absence of its intracellular domains, WFLΔJxK-GFP is unable to interact with the binding partner that is driving WFL polarization to the inner polar domain and the RHID. However, this explanation is not very satisfying as it implies that without its correct binding partner WFL would have a reciprocal localization pattern in H cells (identical to WFLΔJxK-GFP) or that truncated WFL interacts with a different protein that happens to have the opposite polar localization. Further research will be necessary to identify WFL binding partners and determine whether they influence WFL polarity.

The identification of receptor kinases with polar localization provides a new set of conceptual and molecular tools to investigate cell polarity in plants. Unlike transporters, polar localization of receptor proteins is not implicitly tied to their molecular function, suggesting that establishment of polarized signaling domains to perceive extracellular cues is functionally important. There is a long-standing hypothesis that directional signaling and positional information are key drivers of plant development and the identification of polarized receptor kinases, like WFL, supports this hypothesis.

## MATERIALS & METHODS

### Lead contact and materials availability

Further information and requests for resources and reagents should be directed to and will be fulfilled by the Lead Contact, Jaimie Van Norman. Plasmids and transgenic Arabidopsis lines generated in this study have been deposited to the Arabidopsis Resource Center (ABRC, https://abrc.osu.edu/)

### Plant materials and growth conditions

The *Arabidopsis thaliana* Columbia-0 accession was used as the wild type. Standard growth media consisted of 0.5x, 1x, or 1x -Pi Murashige and Skoog (MS) salts (Caisson labs), 0.5 g/L MES (EMD), 1% sucrose, pH 5.7, and 1% agar (Difco), unless otherwise noted. Seeds were surface sterilized with chlorine gas, stratified in tubes at 4°C for 2-3 days and then plated on 100 mm plates with standard growth medium. Plates were then placed vertically in a Percival incubator under long day conditions (16 h light/8 h dark) at a constant temperature of 22°C. Plates were sealed with parafilm for experimental analyses. Seedlings were typically examined between 4-7 days post-stratification (dps). Details for individual experiments are listed in figure legends and/or below.

A candidate insertational allele of *WFL* was obtained from the ABRC (Arabidopsis Resource Center), SAIL_1170_A12, but could not be used for any analyses in this paper as no heterozygous or homozygous mutant individuals could be identified. An allele (*wfl-1*) was generated using CRISPR-Cas9 technology and used for all phenotypic analyses.

### Vector Construction and Plant Transformation

Transcriptional and translational reporter genes were constructed by standard molecular biology methods and utilizing Invitrogen Multisite Gateway® technology (Carlsbad, USA). A region 4.1 kb upstream of the *WFL* (At5g24100) start codon was amplified from Col-0 genomic DNA and recombined into the Invitrogen pENTR™ 5’-TOPO® TA vector. For the transcriptional reporter, the promoter drove endoplasmic reticulum-localized green fluorescent protein (erGFP) as previously described (Van Norman et al., 2014). For translational fusions, the genomic fragment encoding *WFL* from the ATG up to, but excluding the stop codon (including introns, 2.0 kb), was amplified from Col-0 genomic DNA and recombined into the Invitrogen pENTR™ DIRECTIONAL TOPO® (pENTR-D-TOPO) vector and fused to a C-terminal GFP tag (unless otherwise noted) as previously described (Van Norman et al., 2014). Specific primers for *WFL* cloning are listed in Table S1.

WFL-GFP and WFLΔJxK-GFP were driven by cell type-specific promoters (*pSCR*_*2*.*0*_, *pCO*_*2*_, *pC1, pUBQ10*, and *pWER)* as previously described (Lee et al., 2006; Campos et al., 2020). Due to the relatively low fluorescent signal of *pWFL:WFLΔJxK-GFP*, WFLΔJxK-GFP misexpression reporters for these truncations were fused to 3xYFP. *pC1* was received in (Gateway Compatible) pENTR™ P4P1R TA vector from the lab of Philip Benfey, Duke University (Durham, NC, USA). The epidermal translational reporter *pWER:WFL-eYFP:WER3’* was generated as previously described (Campos et al., 2020). The various Gateway compatible fragments were recombined together with the dpGreen-BarT or dpGreen-NorfT destination vector (Lee et al., 2006)).

The dpGreenNorfT was generated by combining the backbone of dpGreenBarT with the p35S::tpCRT1 and terminator insert from pGII0125. Within the target region of the dpGreenBarT, one AclI site was mutated with the QuickChangeXL kit (Stratagene). Plasmids were amplified in ccdB-resistant *E. coli* and plasmids prepped with a Bio Basic Plasmid DNA Miniprep kit. 34uL of the modified dpGreenBarT and unmodified pGII0125 were digested with 1ul each FspI and AclI in CutSmart buffer (NEB) for 1hr at 37C. Digests were subjected to gel electrophoresis on a 1% agarose gel. The 5866bp fragment from the dpGreenBarT and 2592bp fragment from the pGlIII0125 were extracted with a Qiagen MinElute Gel Extraction kit. The fragments were then ligated at 1:1 volumetric ratio (20ng vector; 8.8ng insert) using T4 DNA ligase incubated at 16C overnight before transformation into ccdB-resistant *E. coli*.

Expression constructs were then transformed into Col-0 plants by the floral dip method (Clough and Bent, 1998) using Agrobacterium strain GV3101 (Koncz et al., 1992) and transformants were identified using standard methods. For each reporter gene, T2 lines with a 3:1 ratio of resistant to sensitive seedlings, indicating the transgene is inherited as a single locus, were selected for propagation. These T2 plants were allowed to self and among the subsequent T3 progeny, those with 100% resistant seedlings, indicating that the transgene was homozygous, were used in further analyses. For each reporter, at least three independent lines with the same relative expression levels and localization pattern were selected for imaging by confocal microscopy.

CRISPR-induced mutagenesis was performed as described in (Fauser et al., 2014), a single guide RNA (5’-TTTAACGGTAGTATTCCCGCGGG) was selected in exon 2 of *WFL*. T2 lines that exhibited a 3:1 ratio of resistant to sensitive seedlings, indicating the CRISPR-guideRNA-containing transgene was inherited as a single locus, were selected for continued analyses and sensitive plants were transferred to 1X MS standard growth media to recover. These plants were subsequently tested for lesions in *WFL* in proximity to the guideRNA binding site. We identified *wfl-1*, which has reduced *WFL* expression (Figure S3) and has a single T insertion in the coding region of the second exon that results in a premature stop codon before the transmembrane domain.

### Confocal Microscopy and Image Analysis

Roots were stained with ∼10 μM propidium iodide (PI) solubilized in water for 1-2 min. Imaging was performed via laser scanning confocal microscopy on a Leica SP8 upright microscope equipped with a water-corrected 40x objective and housed in the Van Norman lab. Root meristems were visualized in the median longitudinal or transverse planes. Images were generated using PMT and HYD detectors with the pinholes adjusted to 1 airy unit for each wavelength and system settings were as follows: GFP (excitation 488 nm, emission 492-530 nm), YFP (excitation 514 nm, emission 515-550 nm) and PI (excitation 536 nm, emission 585-660 nm). Unless otherwise indicated, all confocal images are either median longitudinal of roots or transverse sections acquired in the meristematic, elongation, and/or differentiation zones. All plants used for reporter expression imaging were grown on 1x MS with the exception of the roots expressing *pWFL:WFL-GFP* in Figure 1 and Figure 3C and 3E which were grown on 1x MS for 6 days and then transferred to 1x MS -Pi plates for 24 hours. Localization of WFL-GFP was unaffected by -Pi treatment.

For GFP fluorescence intensity measurements of *pWFL:WFL-GFP* and *pWFL:WFLΔJxK-GFP*, seedlings were grown side-by-side on 0.5x MS plates until 5 dps. RHIDsand bulges were selected for analysis at the beginning of the differentiation zone. GFP intensity was measured using Leica (LAS X) quantification software at 3 different positions across the outer epidermal edge. The positions were assigned as follows: Position 1, the area just above the RHID or bulge; Position 2, at center of the RHID or bulge; Position 3, the area just below the RHID or bulge. The maximum GFP intensity was recorded for all 3 positions in 2 biological replicates for 15-20 roots per replicate and GFP intensity was measured in 2-3 cells per root. Representative images of RHID sites and bulges were chosen for each genotype for the figure.

To visualize the dynamic movement of WFL-GFP in RHs, a movie was created by acquiring images every ∼10 seconds for ∼3 minutes. The images were then exported and compiled into a hyperstack using ImageJ software (https://imajej.nih.gov/ij/). These final stacks were saved as an AVI movie with a frame rate of 5 frames per second.

### Phenotypic Analyses

Root hair bulge position measurement protocol was modified from that was previously described in (Masucci and Schiefelbein, 1994b). For root hair bulge position quantification ∼15 roots of each genotype were grown side-by-side on 0.5x MS plates until 4 dps. To image roots, seedlings were stained with PI (as described above) and imaged using confocal microscopy. For each biological replicate, imaging was primarily done in median longitudinal sections at the beginning of the differentiation zone of 15 roots with 3-5 epidermal cells with a root hair bulge selected from each root for analysis. Root hair bulges were binned into 2 categories with the following parameters: “normal (WT)” if there was a measurable distance from the emerging root hair to the rootward edge of the cell and “shifted” if there was no measurable distance. Images were analyzed using ImageJ software for at least 2-4 biological replicates analyzed for each genotype.

### Chemical Treatments

Treatments with small molecules were performed using *pWFL:WFL-GFP* in the Col-0 background seedlings grown on 0.5x MS plates grown until 5 dps. Seedlings were then incubated in liquid 0.5x MS containing one or a combination of the following chemicals: Brefeldin A (BFA) (Sigma-Aldrich) was dissolved in dimethyl sulfoxide (DMSO) (Calbiochem, Cat #317275) in 50 mM stocks and added to the media at a final concentration of 50 µM for 1 hour or other indicated times. Cycloheximide (CHX) (Sigma-Aldrich, Cat #C7698) was added from a 50 mM aqueous stock to a final concentration of 50 µM for 2 hours or other indicated time. Wortmannin (Wm) (Sigma-Aldrich) was dissolved in DMSO and used at 33 µM for 2 hours. In control (mock) experiments, seedlings were incubated in the same media containing an equal amount (0.05% to 0.1%) of the correspondent solvent.

### RT-qPCR Analysis

Total RNA for quantitative RT-PCR (qRT-PCR) was isolated using Qiagen’s RNeasy Plant Mini Kit. Total RNA was extracted from whole seedlings at 7 dps after growth on our standard 1X MS (*wfl-1* and Col-0) or 0.5x MS (Col-0 and *pWFL:WFL-GFP)* growth medium and sealed with parafilm. For each of the biological replicates Col-0 and *wfl-1* or *pWFL:WFL-GFP* were grown side-by-side on the same plate. RNA was isolated for three independent biological replicates for Col-0, *pWFL:WFL-GFP*, and the *wfl-1* allele. First-strand cDNA was synthesized from 1 μg total RNA with RevertAid First Strand cDNA Synthesis and the oligo(dT)_18_ primer (Thermo Scientific). qRT-PCR reactions were set up using IQ SYBR Green Supermix (BioRad) and analysis was performed on the CFX-Connect Real-Time System housed in the Integrative Institute of Genome Biology Genomics Core facility at UC-Riverside. The reaction conditions for each primer pair were: 95°C for 3 min followed by 40 cycles of 95°C for 10s and 57°C for 20 s. Standard curves were performed at least in duplicate. Primer pair efficiency values were calculated for each replicate of the standard curves and the average efficiency was used for subsequent analysis (Table S2). For each genotype and biological replicate, three technical replicates were performed. Data analysis was performed with the Bio-Rad CFX Manager software 3.1 and transcript levels were normalized to *SERINE/THREONINE PROTEIN PHOSPHATASE2A* (*PP2A) (Czechowski et al*., *2005)*.

### Quantification and Statistical Analysis

The Leica LAS X software, as well as ImageJ were used for post-acquisition confocal image processing. The corrected plasma membrane fluorescence intensity data was obtained by analyzing the images with the software ImageJ and calculating the Integrated Density of plasma membrane fluorescence and subtracting (Area of selected cell x Mean fluorescence of background readings). Graphs were generated using PRISM8 (GraphPad Software, https://www.graphpad.com/, San Diego, USA). The exact value of n, what n represents, the number of biological or technical replicates, the means, standard error of the mean (SEM), standard deviation (SD), and how statistical significance was defined are indicated in each of the relevant figure legends. Standard two-tailed student’s t test was performed when comparing wild type to mutant and overexpression phenotypic aspects as a normal distribution is expected.

## ACKNOWLEDGMENTS

We thank members of the Van Norman Lab: Roya Campos and Jason Goff and Dr. Carolyn Rasmussen and Dr. Patricia Springer (UC, Riverside) for discussions of the project and feedback on the manuscript while it was in preparation. We also thank Dr. Erin Sparks (Delaware Biotechnology Institute) for providing the NorfT version of the dpGreen Gateway compatible destination vector. We appreciate access to and assistance from the Institute of Integrative Genome Biology Genomics Core Facility (UC, Riverside). This work was supported by Initial Complement (IC) funds from the University of California, Riverside, USDA-NIFA-CA-R-BPS-5156-H, NSF CAREER award #1751385 to J.M.V.N., and NSF-GRFP award #DGE-1326120 to J.N.T.

## AUTHOR CONTRIBUTIONS

Conceptualization: J.M.V.N.; Methodology and Investigation: J.M.V.N., J.N.T., and C.R-F.; Resources: J.M.V.N. and J.N.T.; Writing - Original Draft: J.N.T.; Writing - Review and Editing, J.N.T., C.R-F., and J.M.V.N.; Visualization: J.N.T. and C.R-F.; Supervision: J.M.V.N.; Funding Acquisition: J.M.V.N. and J.N.T.

## FIGURE LEGENDS

**Supplemental Movie SM1. WFL is a highly dynamic transmembrane protein**. WFL-GFP is dynamic and moves to and from on the plasma membrane during RH development.

---

**Supplemental Figure S1.**
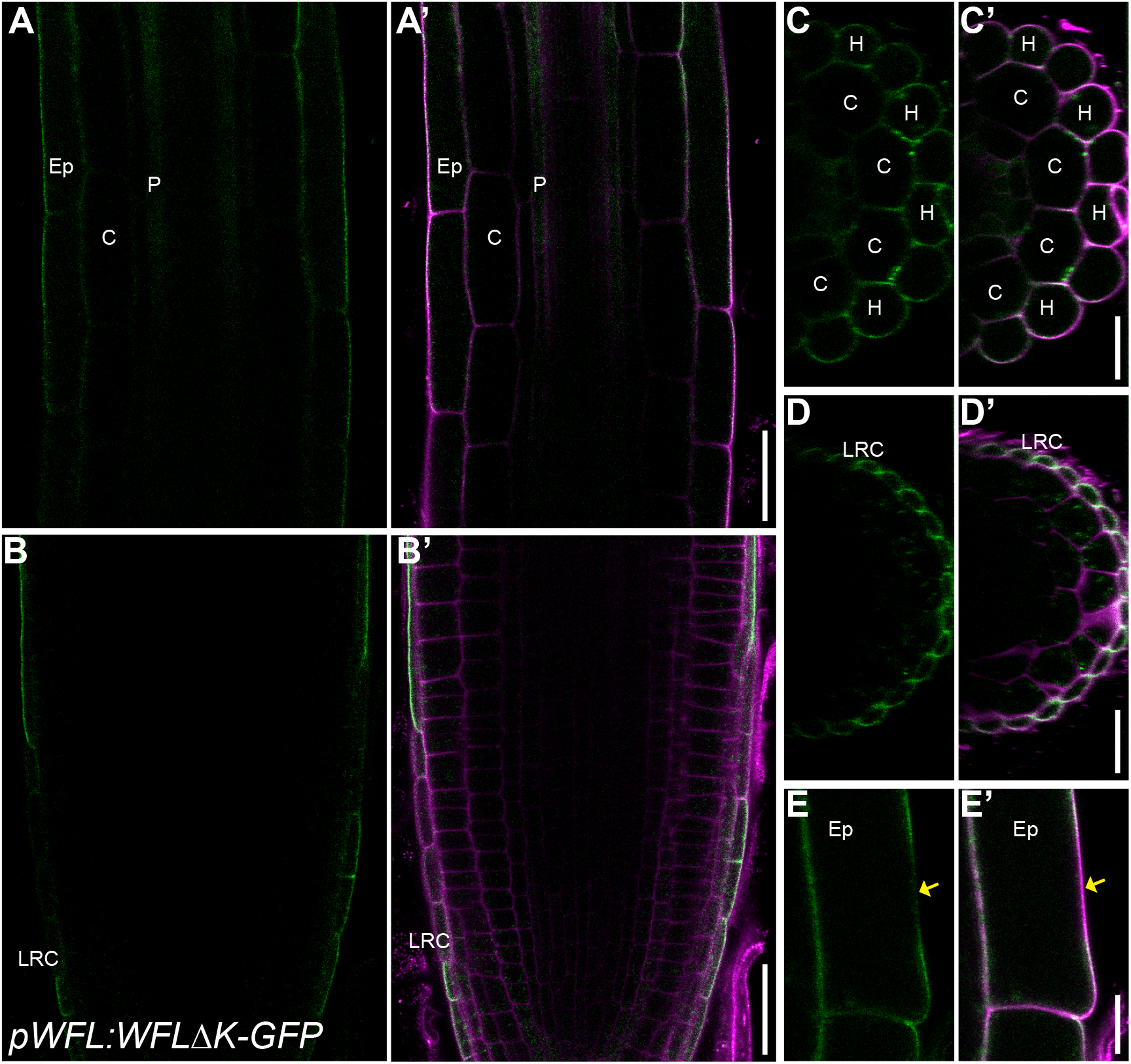
WFLΔK-GFP localizes to the outer polar domain of lateral root cap and epidermal cells. (A-E) Confocal images of WT roots expressing WFLΔK-GFP driven by *pWFL* (*pWFL:WFL*ΔK-GFP) and stained with propidium iodide (PI) to show cell outlines. Adjacent panels show GFP alone (α) and GFP + PI merged (α’). (A and C) In elongation and differentiation zones, WFLΔK-GFP localizes to the outer polar domain of epidermal cells with (C) preferential accumulation in H cells. (E) WFLΔK-GFP is excluded from RHIDs. (B and D) WFLΔK-GFP localizes to the outer polar domain of the LRC. Abbreviations: LRC, lateral root cap; Ep, epidermis; C, cortex; H, hair cell. Scale bars: 25 μm in (A, B, and E); 10 μm in all others.

**Supplemental Figure S2.**
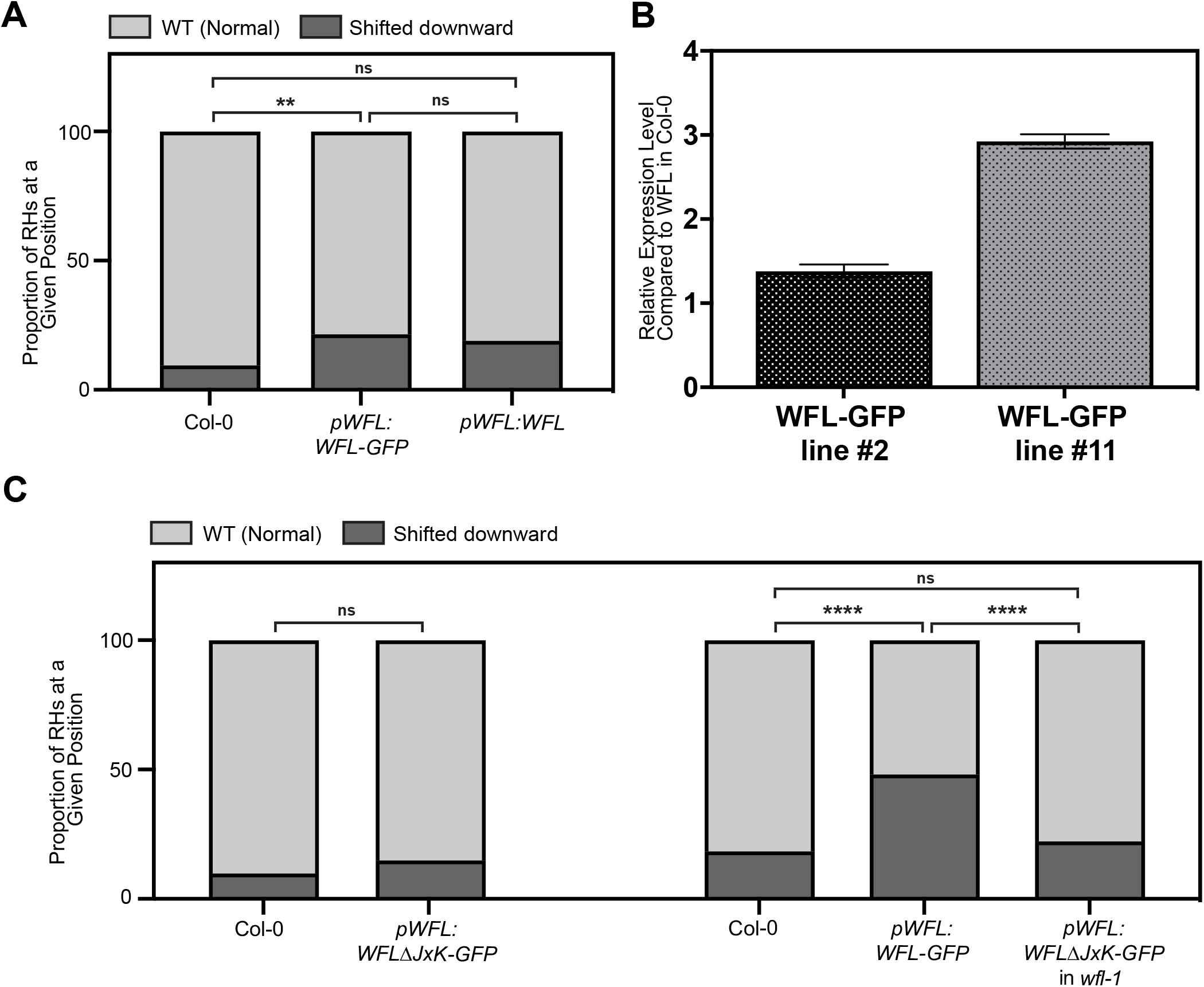
GFP does not cause shifted RH bulge phenotype and this phenotype is not observed in roots expressing *pWFL:WFLJxK-GFP*. (A and C) RH bulges were binned into two categories and quantified. (A) RH bulges are shifted towards the rootward edge of H cells in roots expressing an untagged version of WFL (*pWFL:WFL*). (C) Bulge position is unaffected in roots expressing *pWFL:WFLΔJxK-GFP* and *pWFL:WFLΔJxK-GFP* in *wfl-1*. (B) RT-qPCR showing transcript levels of *WFL* among transgenic lines used for phenotyping. Error bars show standard error of the mean. For RH bulge position graphs: student’s t test, ** p<0.01 and **** p<0.0001. Data shown is from one (of two) independent transgenic lines per reporter (with similar results for each line in each replicate) and 2-3 biological replicates combined, with n= 15 roots and 3-5 cells per root for each replicate.

**Supplemental Figure S3.**
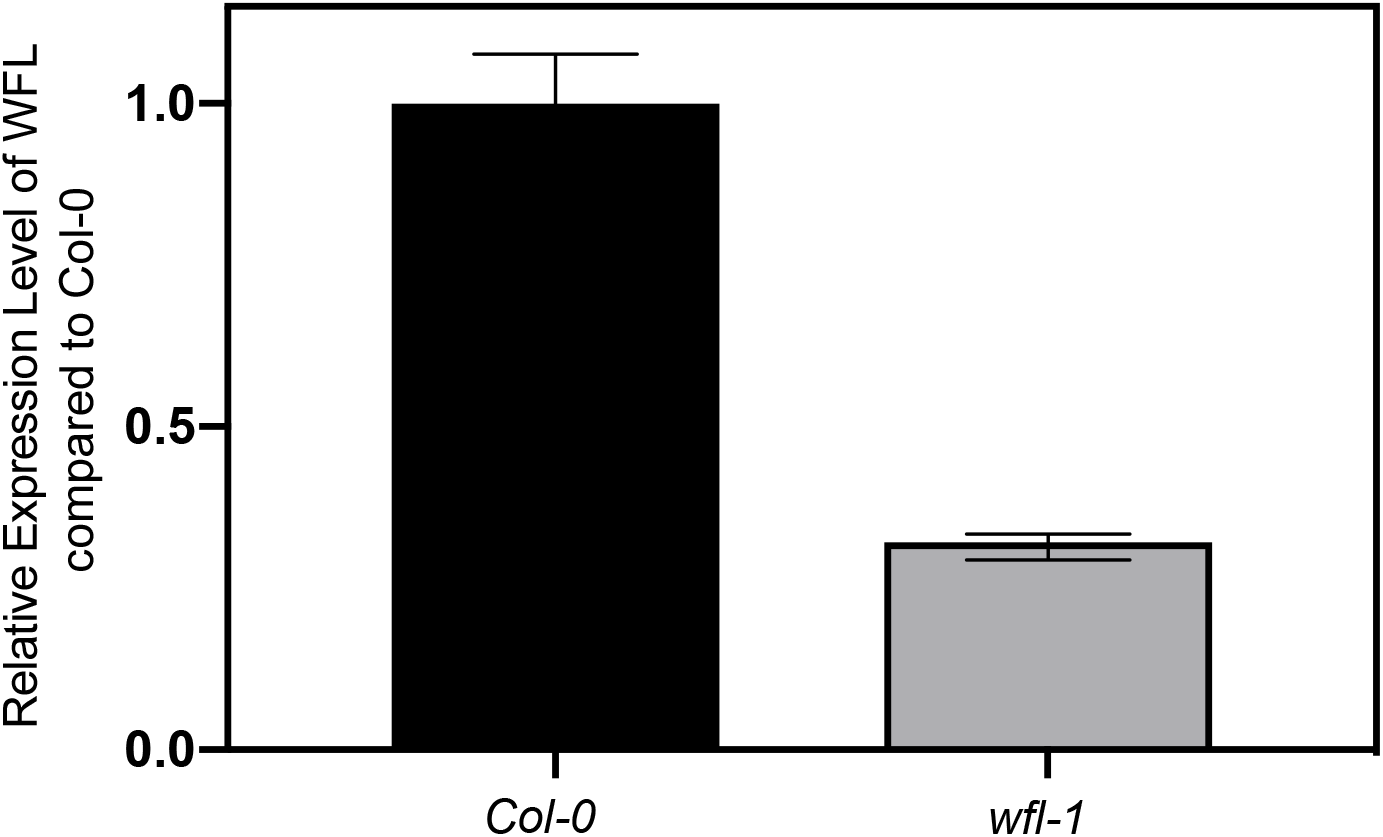
*WFL* transcript level is reduced in *wfl-1*. RT-qPCR showed reduced *WFL* transcript levels in wfl-1. *WFL* expression is relative to *SERINE/THREONINE PROTEIN PHOSPHATASE 2A* (*PP2A*). Data shown for one biological replicate (of three) with 3 technical replicates performed per experiment and each experiment was repeated 3 times. Error bars indicate standard error of the mean.

**Table S1.**
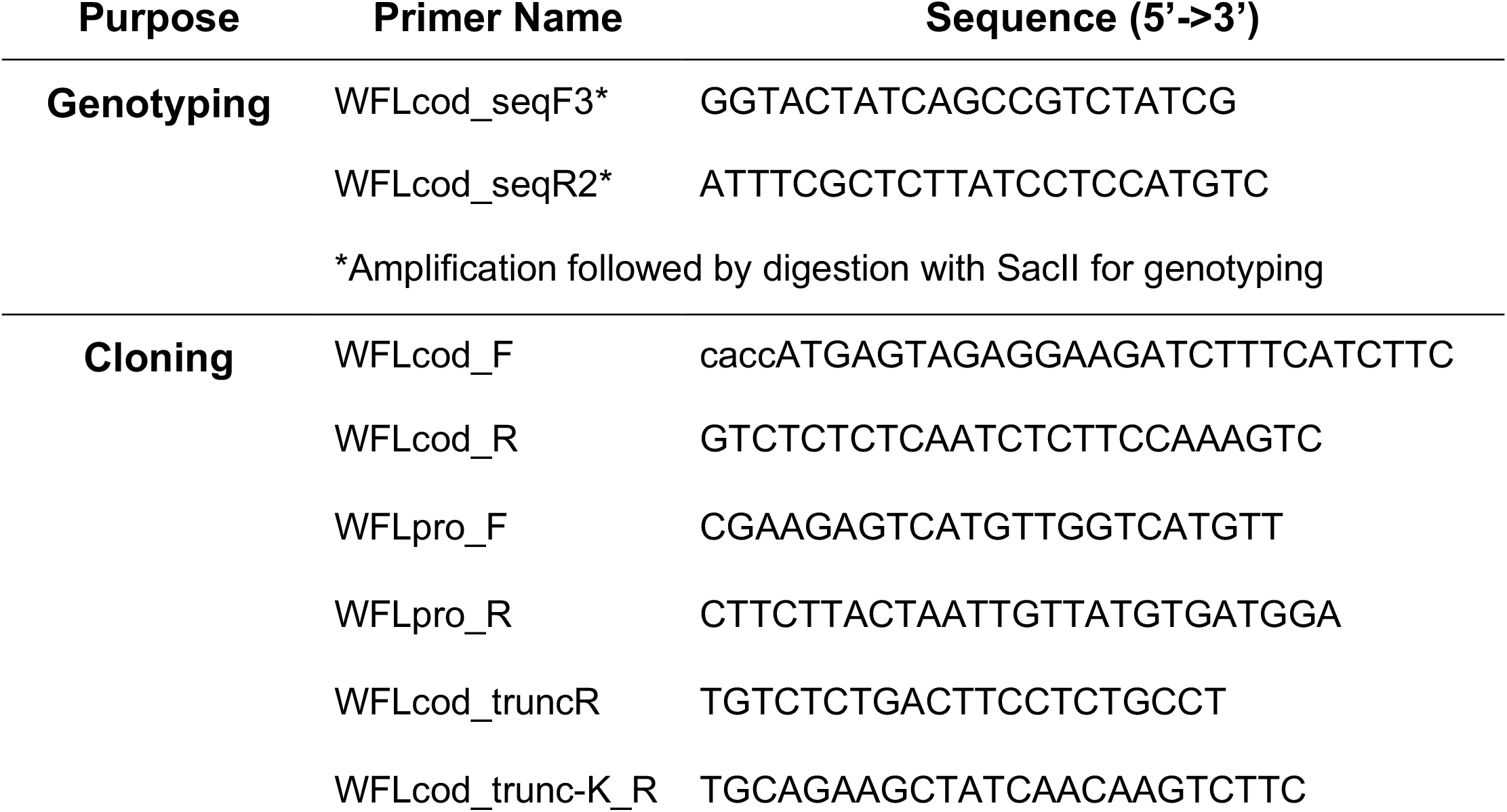
Cloning and genotyping primers.

**Table S2.**
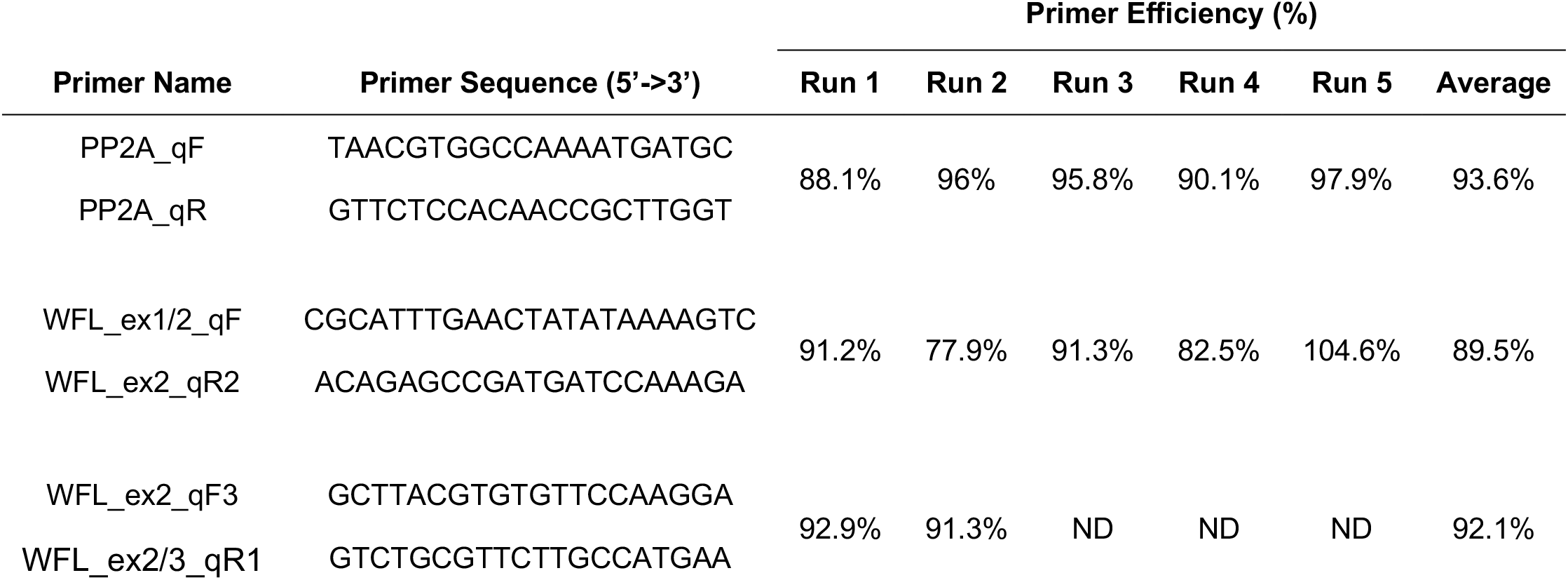
Primers and primer efficiency information for RT-qPCR.

